# Knockdown of TTLL1 reduces Aβ-induced TAU pathology in human iPSC-derived cortical neurons

**DOI:** 10.1101/2024.11.19.624324

**Authors:** Mohamed Aghyad Al Kabbani, Laura Köhler, Tamara Wied, Daniel Adam, Jennifer Klimek, Hans Zempel

**Author notes:** Correspondence to: Dr. Dr. Hans Zempel.

## Abstract

Microtubules play a crucial role in neuronal structure and function, with their stability and dynamics regulated by posttranslational modifications (PTMs) such as polyglutamylation. In Alzheimer disease (AD), the microtubule-associated protein TAU becomes mislocalized into the somatodendritic compartment (‘TAU missorting’), dissociates from microtubules, aggregates into neurofibrillary tangles, and contributes to microtubule destabilization and neuronal death. Here, we investigated the role of Tubulin-Tyrosine-Ligase-Like proteins (TTLLs) in TAU missorting and microtubule dysregulation using human induced pluripotent stem cell (hiPSC)-derived cortical neurons treated with oligomeric amyloid-beta (oAβ) to replicate AD-like conditions. TTLL1, TTLL4, TTLL6 were selectively knocked down (KD) to assess their impact on TAU missorting and microtubule stability. Fluorescence resonance energy transfer (FRET) microscopy was used to examine interactions between TAU and TTLL proteins. We observed TAU missorting, increased tubulin polyglutamylation, decreased microtubule stability, and synaptic declustering in oAβ-treated neurons. TTLL1 KD significantly reduced TAU missorting, tubulin polyglutamylation, and synaptic disintegration, while TTLL4 KD showed moderate effects, and TTLL6 KD restored microtubule acetylation. Importantly, TTLL KD did not impair neuritic networks, dendritic complexity, or neuronal activity. FRET microscopy revealed a potential interaction between TAU and TTLL1, but not other TTLLs, suggesting a direct role of TTLL1 in TAU-mediated toxicity. Our findings indicate that targeting TTLL1, either alone or in combination with other TTLLs, may be a promising therapeutic strategy to counteract microtubule and synaptic dysfunction in AD and related neurodegenerative disorders.

## Introduction

Microtubules are cylindrical filamentous heterodimers of α- and β-tubulin that form an essential part of the cytoskeleton. Microtubules play an important role in maintaining neuronal shape and facilitating the transportation of organelles and vesicles across neuronal networks (Sakakibara et al., 2013). Microtubule dynamics and functions are tightly regulated by a complex set of posttranslational modifications (PTMs) (Janke and Kneussel, 2010), as well as binding to and interacting with proteins like microtubule-associated proteins (MAPs) and microtubule-severing enzymes such as spastin (Goodson and Jonasson, 2018). In Alzheimer disease (AD), microtubules are significantly depleted in affected brains, though the underlying causes remain unclear (Cash et al., 2003; Jean and Bass, 2013).

TAU, a microtubule-associated protein encoded by the *MAPT* gene, is an axonal-enriched protein that binds microtubules and promotes their assembly and stability. However, in tauopathies like AD, TAU dissociates from microtubules, missorts to the somatodendritic compartments, and aggregates into hyperphosphorylated neurofibrillary tangles, leading to microtubule fragmentation and neuronal death (Zempel and Mandelkow, 2015). Spastin, an ATP-dependent enzyme encoded by the *SPAST* gene, mediates microtubule severing, and its activity is catalyzed by a specific PTM of microtubules called polyglutamylation, which comprises the addition of a glutamate side chain to a glutamate residue, usually on the C-terminal tail of tubulin (Lacroix et al., 2010). This modification is carried out by several members of a class of enzymes known as Tubulin-Tyrosine-Ligase-Like proteins (TTLLs), mainly TTLL 1, 4, 5, 6, 7 and 11 (Magiera et al., 2018).

Polyglutamylation is reversible, with cytosolic carboxypeptidases (CCPs) removing glutamate side chains (Rogowski et al., 2010). The balance between adding and removing glutamate chains is essential for health, with hyperglutamylation linked to neurodegeneration. For instance, Purkinje cell degeneration (pcd) mice, in which CCP1 is deficient, exhibit severe neurodegeneration, which is attenuated by knocking out TTLL1 and TTLL4, but not TTLL5 or TTLL7 (Li et al., 2020; Wu et al., 2022). Another hint towards potential involvement of TTLLs in neurodegeneration came from primary rat neurons treated with oligomeric amyloid-beta (oAβ), where missorted TAU appeared to recruit TTLL6 to the somatodendritic compartments, inducing polyglutamylation and spastin-mediated microtubule severing, leading to extensive microtubule loss (Zempel et al., 2010, 2013). However, the underlying and potentially druggable disease mechanisms by which TAU missorting triggers microtubule dysfunction, and the specific TTLLs involved, remain unclear in human disease-relevant models.

In this study, we first aimed to establish a human tauopathy-relevant model using human induced pluripotent stem cell (hiPSC)-derived cortical neurons (iNeurons). We treated these neurons with oAβ and observed TAU missorting, increased tubulin polyglutamylation, and synaptic declustering. We then aimed to identify the TTLL(s) responsible for mediating the pathological effects of missorted TAU via individually knocking down TTLL1, TTLL4, or TTLL6. We show that decreased expression of TTLL1, and to some extent TTLL4, alleviate the pathological effects of oAβ without disturbing neuritic network, dendritic morphology, or neuronal function. We provide first evidence that targeting TTLL1 alone or in combination with another TTLL is a potential therapeutic target in AD and other tauopathies, and should be further validated and investigated.

## Methods

### hiPSC maintenance

WTC11 cells with a doxycycline-inducible Ngn2 transgene (Miyaoka et al., 2014; Wang et al., 2017) were cultured on Geltrex-coated plates (Thermofisher Scientific #A1413302) at 37°C, 5% CO_2_ and regularly passaged when almost fully confluent using Versene (Thermofisher Scientific #15040066) and thiazovivin-supplemented StemMACS iPS-Brew X.F. (Axon Medchem #1535, Miltenyi Biotec #130-104-368) for the first 24 hours (Buchholz et al., 2024).

### Differentiation of hiPSCs into cortical neurons (iNeurons)

Differentiation of hiPSCs into cortical neurons was carried out as described before (Buchholz et al., 2024; Bachmann et al., 2021; Wang et al., 2017). At the start of differentiation, iPSCs were harvested using Accutase (Sigma-Aldrich #A6964-100ML) and seeded onto Geltrex-coated plates pre-differentiation medium (Thermofisher Scientific #12660012) supplemented with thiazovivin (day before differentiation: d−3). The medium was changed every day for 2 days to fresh pre-differentiation medium without supplementation. On day 0, 50,000 or 300,000 cells were seeded onto Poly-D-Lysine- (Sigma-Aldrich #P7886-50MG) and Laminin- (Trevigen #3446-005-01) coated 24-well-plates or 6-well plates respectively using maturation medium supplemented with 1:100 GelTrex. Half of the media was exchanged once per week until analysis.

### oAβ preparation and treatment

Aβ was prepared and reconstituted into oligomers as described before (Zempel et al., 2013). Briefly, Aβ40 and Aβ42 powder (rPeptide #A-1153-1 and #A-1163-2) were completely dissolved in Hexafluoro-2-propanol (HFIP) to a final concentration of 1 mM. Following aliquotation, HFIP was evaporated completely using a vacuum concentrator, and the lyophilized powder was stored at −80°C. On the day of treatment of iNeurons, lyophilized Aβ40 and Aβ42 were redissolved in 50 mM NaOH and mixed to produce a Aβ40/Aβ42 ratio of 7:3, and then diluted with PBS and 50 mM HCl to a final concentration of 100 µM. To induce oligomerization, Aβ mixture was incubated at 37°C for one hour. Subsequently, iNeurons at day 21 were treated with 1 µM oAβ for 3 hours and analyzed. Fig. S1 shows oAβ-treated iNeurons highlighting the distribution and targeting of the oligomers.

### Short hairpin RNA sequences

For the knockdown of human *TTLL*s in iNeurons, short hairpin RNA (shRNA) oligonucleotides were inserted into pLKO.3G vector (Addgene #14748) resulting in a multi-cistronic lentiviral construct expressing Green fluorescent protein (GFP) and the corresponding shRNA. The following shRNA sequences were used to target human *TTLL*s or as a control:

Scrambled shRNA (control): 5’-TTGTCTTGCATTCGACTAA-3’

sh*TTLL1*: 5’-GTTTGTGTCTCAATCTAATAA-3’

sh*TTLL4*: 5’-GAGCCTTGGCAATAAGTTC-3’

sh*TTLL6*: 5’-CGGACUCATGAUUUCCAGGATT-3’, 5’-AACAACUCCCUCUUCCAGAAU-3’

### Lentiviral-based knockdown of TTLLs

Lentivirus particle production and subsequent lentiviral transduction of iNeurons are described in detail in Buchholz et al., 2024. Briefly, HEK293T cells were co-transfected with the corresponding pLKO.3G plasmid, the packaging plasmid psPAX, and the envelope plasmid pMD2.G (Addgene #12259 and #12260). Four and five days after transfection, the culture supernatant containing lentivirus was collected, filtered and stored at −80°C. iNeurons were transduced with the lentiviral particles on day 10 and analyzed 11 days after transduction (day 21).

### Western blot analysis

For Western blot analysis, iNeurons were lysed in RIPA buffer (Sigma Aldrich #R0278), centrifuged at 16,000 ×g for 20 minutes at 4°C, diluted in 5× Laemmli buffer, boiled for 10 minutes at 95°C, and then separated on 10% Sodium dodecyl sulfate (SDS)-polyacrylamide gels. Afterwards, proteins were transferred to polyvinylidene fluoride (PVDF) membranes overnight at 4°C, and blocked in 5% bovine serum albumin (BSA) in Tris-buffered saline with 0.1% Tween (TBS-T). Membranes were incubated with the primary antibody overnight at 4°C, washed three times with TBS-T, and incubated with the corresponding secondary horseradish peroxidase (HRP)-coupled antibody for 1 hour at room temperature. After three washing rounds with TBS-T, the immunoreactions were detected by applying the SuperSignal West Pico Chemiluminescent Substrate (Thermofisher Scientific #34580) using a ChemiDoc XRS + system (Bio-Rad).

### Immunofluorescence labeling of iNeurons

For immunocytochemistry, iNeurons were fixed with 3.7% Formaldehyde in PBS containing 4% sucrose at room temperature for 30 minutes. Afterwards, cells were permeabilized and blocked in 5% BSA (Carl Roth #8076.4) and 0.2% Triton X-100 (Carl Roth #3051.2) in PBS for 10 minutes. After fixation, iNeurons were stained with primary antibodies at 4°C overnight. The following day, coverslips were washed three times with PBS and stained with the corresponding secondary antibodies coupled to an AlexaFluor dye for two hours at room temperature. Coverslips were then washed with PBS and stained with NucBlue (Thermofisher Scientific #R37605) for 20 min at room temperature, followed by mounting onto glass slides using Aqua-Poly/Mount (Polysciences #18606-20). The slides were dried 24 hours at room temperature and then imaged.

### Imaging

Immunostained iNeurons were imaged using a wide-field fluorescence microscope (Axioscope 5, Zeiss) with ZenBlue Pro imaging software (V2.5, Zeiss). Images were analyzed using ImageJ software (Version 2.14.0/1.54f, National Institutes of Health and the Laboratory for Optical and Computational Instrumentation (LOCI), University of Wisconsin, USA). To measure the levels of TAU or tubulin PTMs in neurons, regions of interest (ROIs) were manually delineated as profiles of the soma or the dendrite where no other somas or processes overlapped, and the mean fluorescence intensity (MFI) of each ROI was measured.

### Neuronal network analysis

Fields of iNeurons cultures were imaged at 10x magnification after immunostaining for the axon-enriched neurofilament light chain (NF-L) and the somatodendritic marker microtubule-associated protein 2 (MAP2). The area of NF-L- or MAP2-positive neurites was calculated using ImageJ software and normalized to the number of transduced nuclei in each field, identified via GFP fluorescence.

### Sholl analysis

Sholl analysis (Sholl, 1953) was used to investigate the complexity of dendritic arbo uring in iNeurons immunostained for MAP2. The center of the soma was designated as the center of concentric circles, and the number of intersections was analyzed via the Neuroanatomy plugin in ImageJ software. Sholl profiles were created by plotting the number of intersections against the distance from the soma (µm). For statistical comparison, either the area under the curve (AUC) or the full-width at half maximum of the Sholl profiles were compared. Data was tested for normality by Shapiro-Wilk test and compared using a pairwise t-test. Analysis was performed using R (R Core Team, 2024), AUC was calculated using the DescTools package (Signorell, 2024).

### Microelectrode array measurements

For microelectrode array (MEA) measurements, iPSCs were seeded on MEA 24-well plates and differentiated into cortical neurons as described above. At day 10, iNeurons were transduced with lentiviral particles carrying the corresponding shRNA. Spontaneous activity was recorded for 2 minutes at day 21 at 37°C.

### Fluorescence Resonance Energy Transfer (FRET) assay

Live-cell imaging-based FRET assay was carried out to assess interactions between TAU and different TTLLs. HEK293T cells were co-transfected for 24 hours with teal fluorescent protein (TFP)-TAU construct (FRET donor) and yellow fluorescent protein (YFP)-TTLL1, TTLL4, or TTLL6 constructs (FRET acceptor). HEK293T cells were also transfected for 24 hours with empty vectors expressing TFP or YFP as controls. Live-cell imaging was performed with an inverted Leica DMi8 microscope with the help of Leica LAS X software (v3.7.3). FRET efficiency was analyzed via FRET and Colocalization Analyzer plugin (Hachet-Hass et al., 2006) on ImageJ software.

### Statistical analysis

The GraphPad Prism (v9.5.1, GraphPad Software, Boston, Massachusetts, USA) was used for statistical analysis. Shapiro-Wilk test was performed to test for normal distribution of the data. In case of normal distribution, statistical analysis was performed by unpaired t-test to compare the means of two groups, or one-way ANOVA with correction for multiple comparisons (Tukey’s test) to compare three or more groups. When the data were not normally distributed, Mann –Whitney U test or Kruskal-Wallis test with correction for multiple comparisons (Dunn’s test) were carried out, respectively. Statistical significance was denoted by a significance level of P < 0.05.

### Antibodies

The antibodies used in this study are listed in Table 1.

**Table 1.**
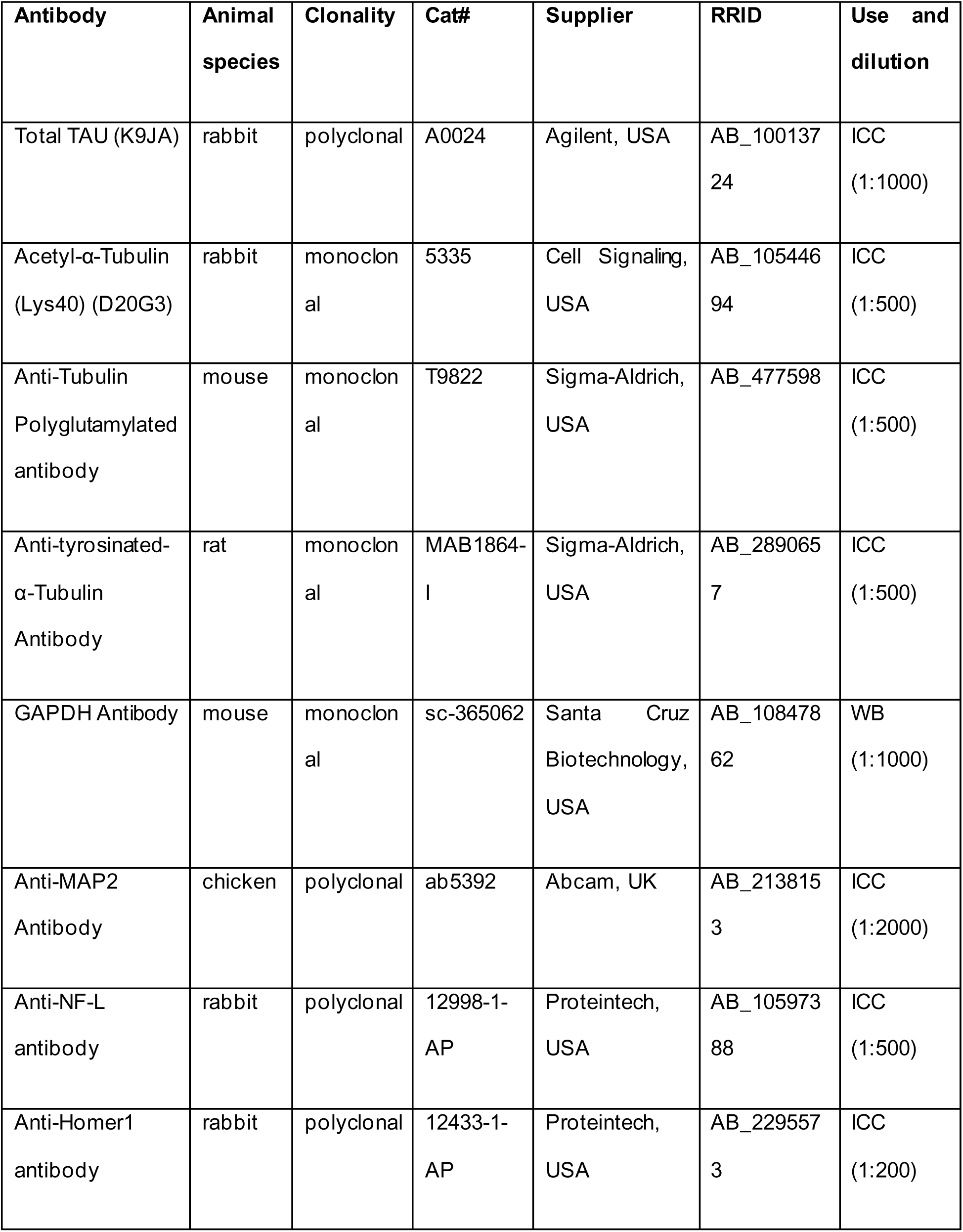
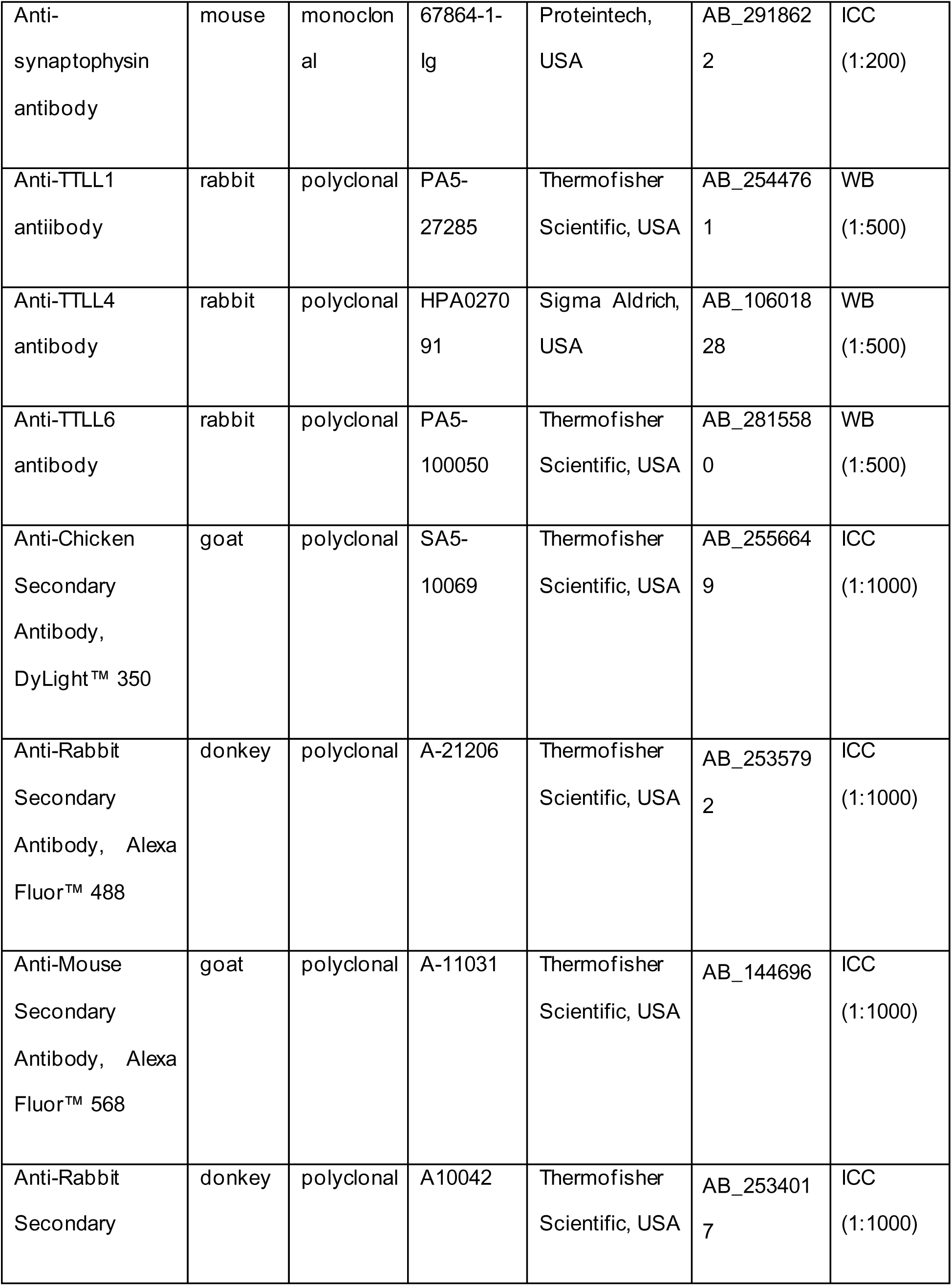
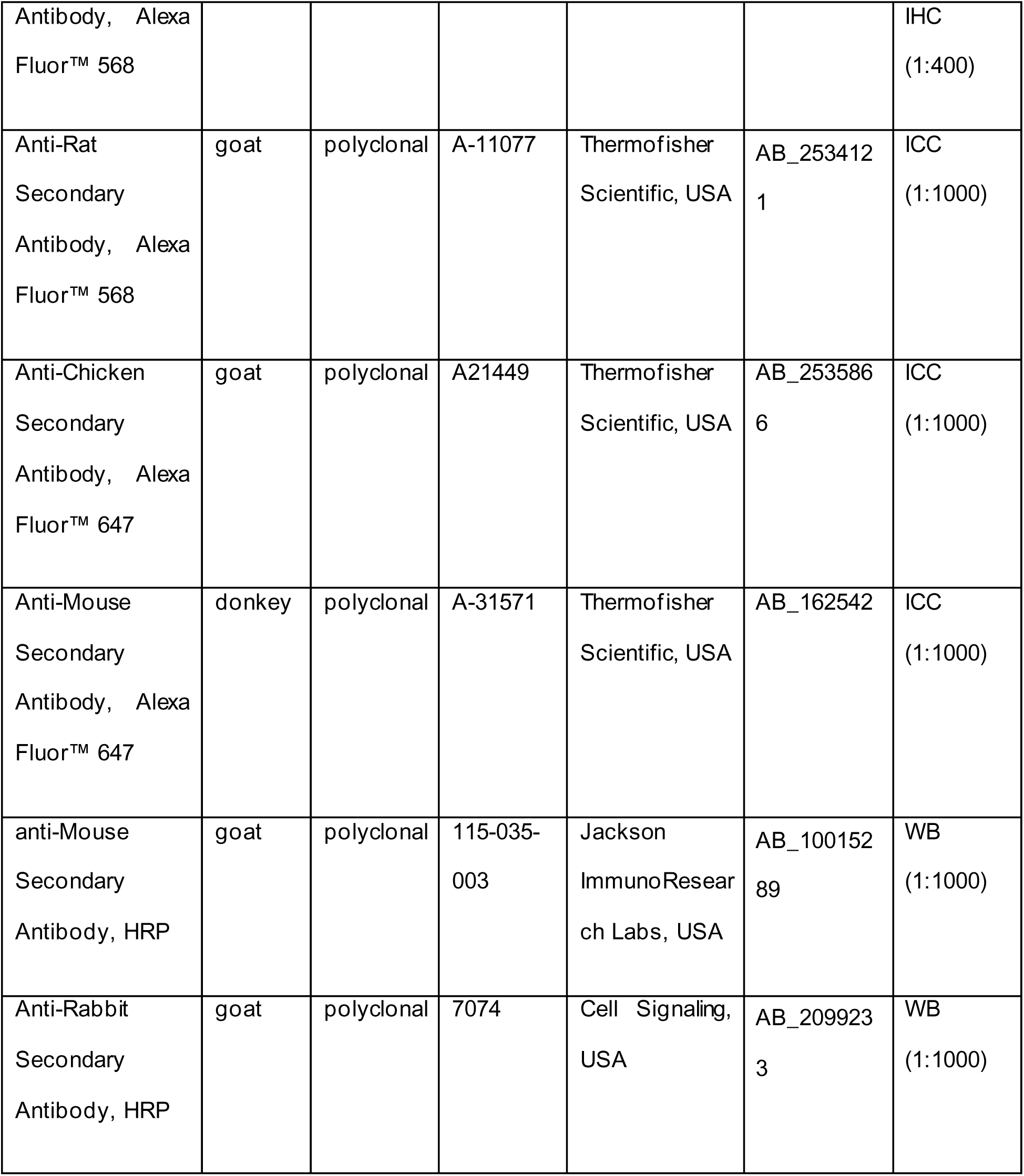
List of the primary and secondary antibodies used in the study.

## Results

### Establishment of Aβ-induced TAU pathology in iNeurons

Previous insights into the potential connection between TAU missorting, microtubule instability, and TTLLs were obtained from rodent-derived models, while a representative human neuronal model was lacking. In order to establish a human tauopathy-relevant model to study TAU-based effects on microtubules, human iPSCs were differentiatedinto cortical neurons. Briefly, genetically modified WTC11 iPSC line harboring a doxycycline-inducible neurogenin-2 (Ngn2) transgene were induced to differentiate into pure glutamatergic neuronal cultures (See: Methods in this paper; Wang et al., 2017; Buchholz et al., 2024).

Day 21 iNeurons treated with 1 μM oAβ showed increased levels of somatic TAU indicative of TAU missorting (Fig. 1A, E). This was accompanied by a marked increase in the levels of tubulin polyglutamylation (Fig. 1B, F). Interestingly, levels of polyglutamylated tubulin remained unchanged in *MAPT* knockout iNeurons (characterized in Buchholz et al., 2025, Fig. S2). To further evaluate the stability of microtubules in oAβ-treated iNeurons, we investigated the levels of acetylated and tyrosinated tubulin, and observed reduced acetylation in the dendrites and decreased tyrosination in the somatic compartments (Fig. 1C, G, H), indicating that microtubule stability and dynamics were further compromised.

**Figure 1.**
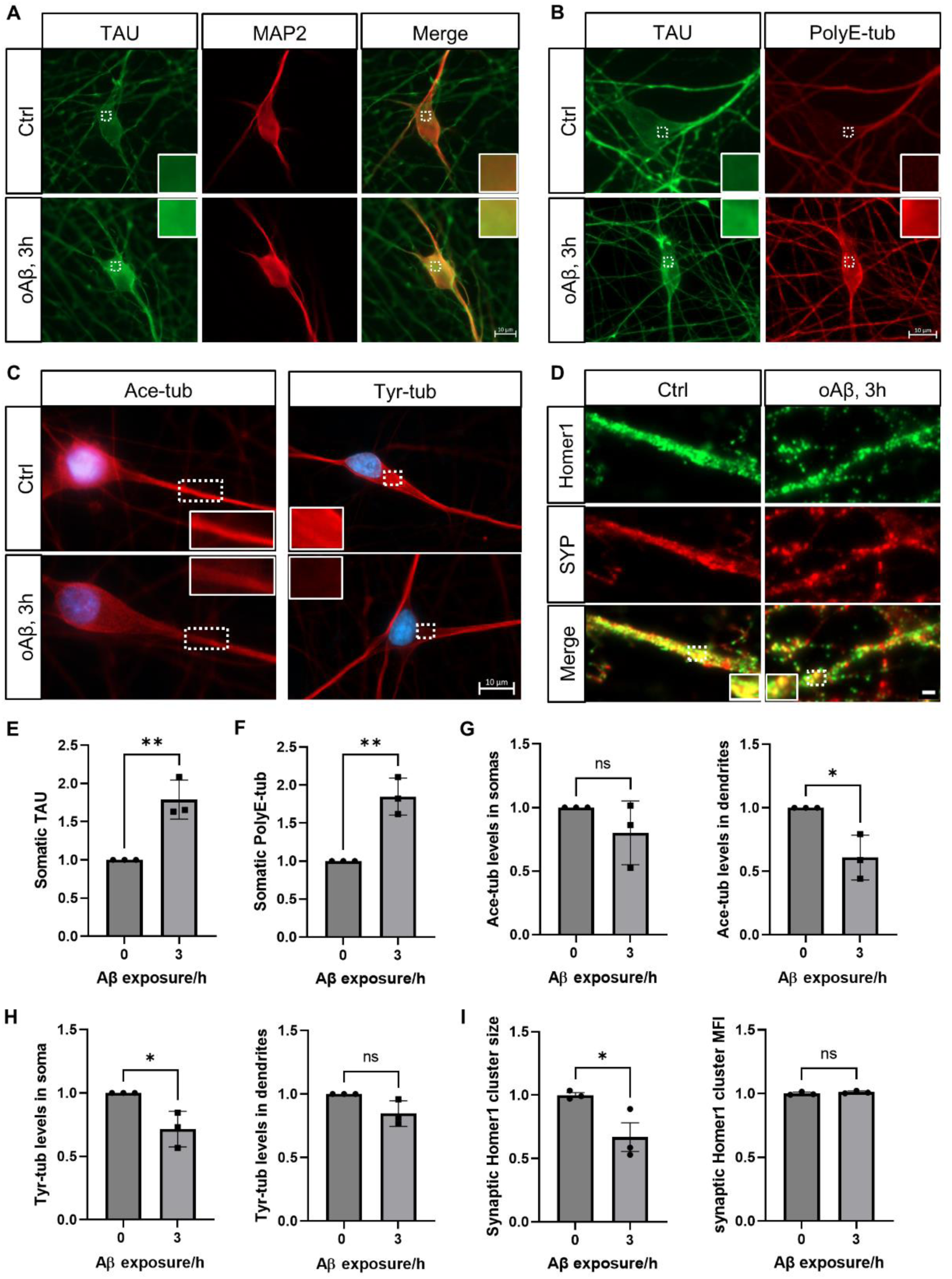
oAβ treatment induces TAU missorting, alters microtubule modifications, and disrupts synaptic clusters in iNeurons. **A)** Co-immunostaining of TAU and MAP2 in iNeurons treated with oAβ for 3 hours or untreated control (Ctrl). Insets display 4-fold magnifications of areas framed with dashed lines, highlighting TAU missorting into the somatic compartment. Scale bar: 10 µm. **B)** Co-staining of TAU and polyglutamylated tubulin (polyE-tub) in iNeurons treated with oAβ for 3 hours or untreated control. Insets show 4-fold magnifications of areas framed with dashed lines, emphasizing increased polyglutamylation in oAβ-treated neurons. Scale bar: 10 µm. **C)** Immunostaining of acetylated tubulin (ace-tub) and tyrosinated tubulin (tyr-tub) in iNeurons treated with oAβ for 3 hours or untreated control. Insets show 3-fold magnifications of areas framed with dashed lines, demonstrating reduced acetylation and tyrosination post-treatment. Scale bar: 10 µm. **D)** Co-immunostaining of Homer1 and synaptophysin (SYP) in iNeurons treated with oAβ for 3 hours or untreated control. Insets show 2-fold magnifications of the areas framed with dashed lines, illustrating the disassembly of synaptic clusters. Scale bar: 2 µm. **E)** Quantification of somatic TAU levels in (A). (N=3, n=45 neurons). **F)** Quantification of somatic polyE-tub levels in (B). (N=3, n=45 neurons). **G)** Quantification of somatic and dendritic ace-tub levels in (C). (N=3, n=45 neurons). **H)** Quantification of somatic and dendritic tyr-tub levels in (A). (N=3, n=45 neurons). **I)** Quantification of SYP-colocalizing Homer1 cluster size and mean fluorescence intensity (MFI) in (D). (N=3, n=150 puncta). ^ns^ non-significance, * P ≤ 0.05, ** P ≤ 0.01.

We also wanted to test the effects of oAβ on the synaptic integrity in our iNeurons. To this end, we triple stained Homer1, synaptophysin, and MAP2 to identify dendritic synapses and quantified the size and mean fluorescence intensity of synaptophysin-colocalized Homer1. While the fluorescence intensity of synaptic Homer1 clusters did not change, the size of these clusters was reduced upon oAβ treatment, indicating synaptic destabilization (Fig. 1D, I). Hence, oAβ treatment in iNeurons causes TAU missorting, microtubule instability, and synaptic instability, making these neurons a suitable human model for our study.

### Reduction of TTLL1 and TTLL4 expression attenuates oAβ toxicity and TAU missorting

To identify the specific TTLL(s) driving pathological microtubule polyglutamylation upon TAU missorting, we aimed to knock down three polyglutamylating TTLLs and then to observe the effects of subsequent oAβ insult on the levels of polyglutamylated tubulin, and on the other toxicity read-outs established above.

TTLL1, TTLL4, and TTLL6 were individually knocked down using shRNA-lentiviral transduction of Day 10 iNeurons. Effective knockdown was confirmed via western blotting at Day 21, with residual expression of targeted TTLLs reduced to 20-50% of the control (iNeurons transduced with viruses carrying scrambled shRNA) (Fig. 2B).

**Figure 2.**
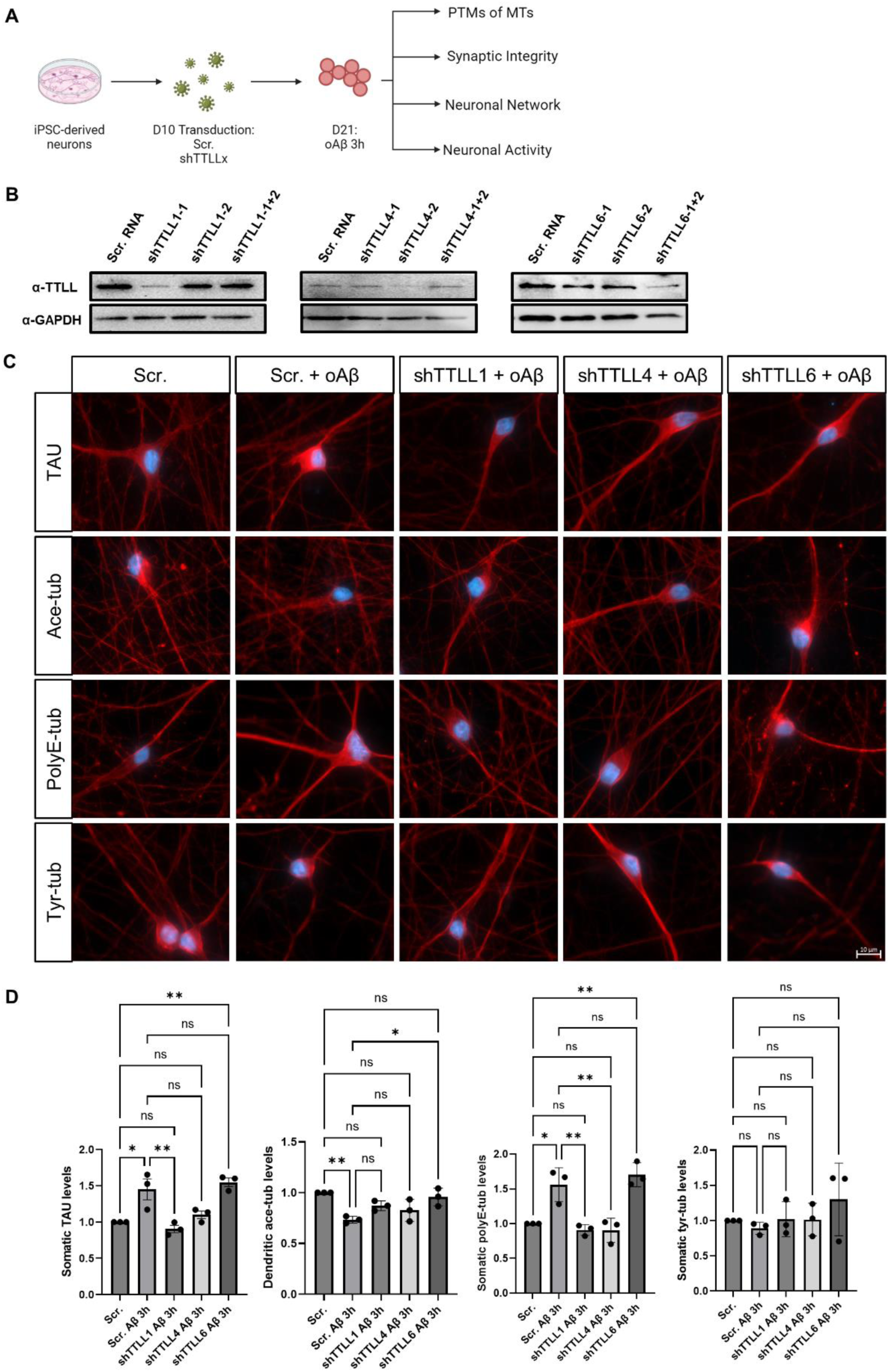
Knockdown of TTLL1 and TTLL4 partially ameliorates oAβ-induced pathological changes in iNeurons. **A)** Schematic representation of the experimental workflow, outlining the knockdown and treatment procedures. Neurons were transduced at Day 10 with shRNA, and then treated and fixed as indicated. **B)** Immunoblotting confirmation of successful knockdown of TTLL1, TTLL4, and TTLL6 in iNeurons. **C)** Immunostaining for TAU, acetylated tubulin (Ace-tub), polyglutamylated tubulin (PolyE-tub), tyrosinated tubulin (Tyr-tub) in Day 21 iNeurons transduced with scrambled RNA (Scr.) or shRNAs targeting TTLL1, TTLL4, or TTLL6 for 11 days, and treated with oAβ or vehicle control for 3 hours. Scale bar: 10 µm. **D)** Quantification of somatic TAU, dendritic ace-tub, somatic polyE-tub, somatic tyr-tub levels of (C). (N=3, n=45 neurons). ^ns^ non-significance, * P ≤ 0.05, ** P ≤ 0.01.

Following the establishment of efficient lentiviral-based knockdown of TTLL1, TTLL4, and TTLL6, the effects of each of these knockdowns on oAβ-induced toxicity was investigated. Day10 iNeurons were transduced with the knockdown viruses or scrambled control. At Day 21, transduced iNeurons were treated with either oAβ or a vehicle control for 3 hours, followed by fixation and staining as above. (Fig. 2A). Interestingly, oAβ-induced TAU missorting and tubulin polyglutamylated tubulin were significantly reduced by TTLL1 knockdown to near normal levels, with TTLL4 knockdown showing a similar effect on polyglutamylation but only a partial effect on TAU missorting. On the other hand, TTLL6 knockdown significantly restored acetylated tubulin levels but did not impact TAU missorting or tubulin polyglutamylation. However, the knockdown of none of these TTLLs was sufficient to counteract the decrease of tyrosinated tubulin induced by oAβ treatment (Fig. 2C-D).

While neither TTLL1 nor TTLL4 knockdowns fully restored synaptic Homer1 cluster size, they both prevented significant cluster disassembly by 15-30 % upon oAβ insult, a protective effect not observed with TTLL6 knockdown (Fig. 3A-B). This indicates that knocking down TTLL1 and TTLL4 in iNeurons reduced oAβ-induced TAU missorting, tubulin polyglutamylation, and partially protected synapses, while TTLL6 restored acetylated tubulin but did not affect TAU missorting or synaptic protection, with none preventing tyrosinated tubulin loss.

**Figure 3.**
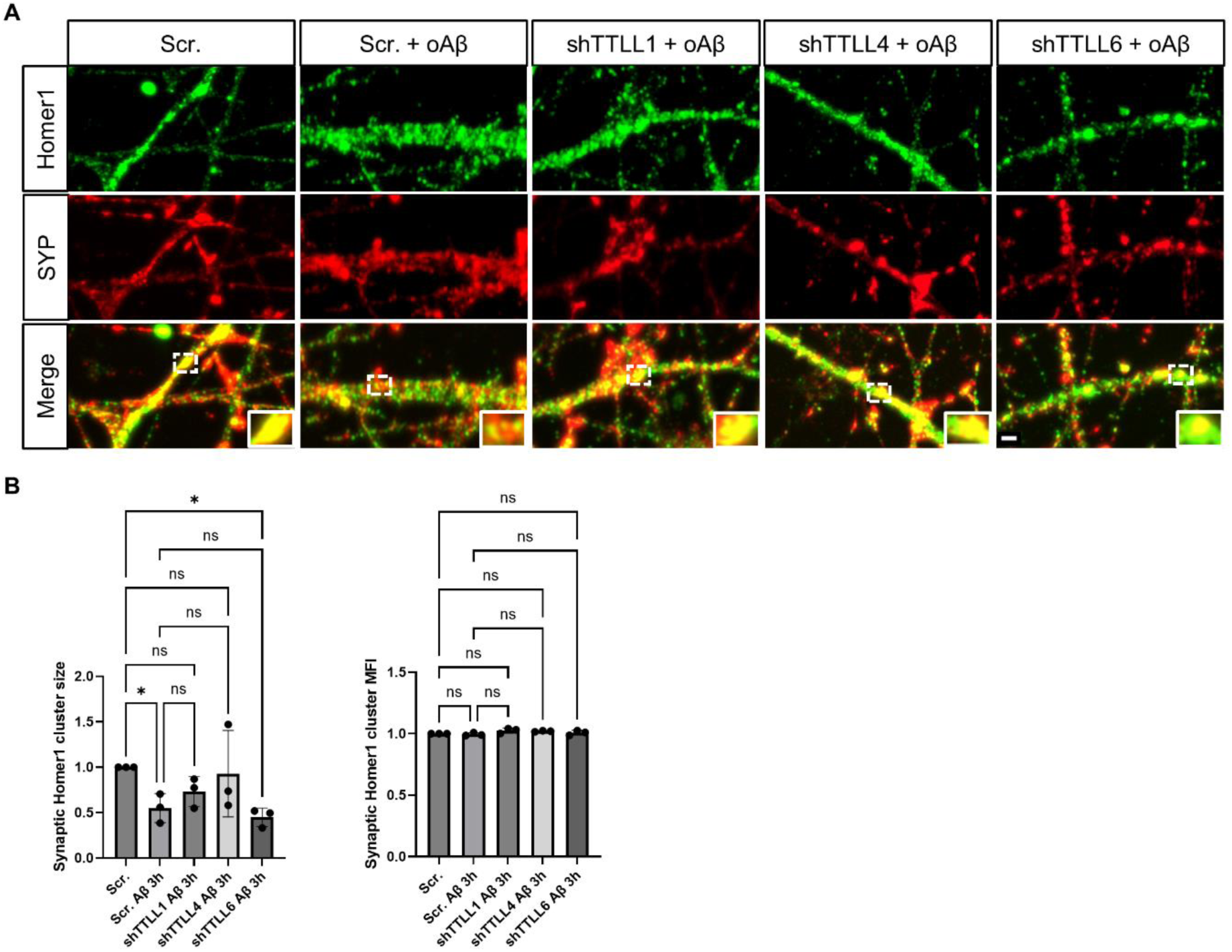
Knockdown of TTLL1 and TTLL4 sightly attenuates oAβ-induced synaptic declustering in iNeurons. **A)** Co-immunostaining of Homer1 and synaptophysin (SYP) in Day 21 iNeurons transduced with scrambled RNA (Scr.) or shRNAs targeting TTLL1, TTLL4, or TTLL6 for 11 days, and treated with oAβ or vehicle control for 3 hours. Insets show 2-fold magnifications of areas framed with dashed lines, highlighting changes in synaptic cluster size. Scale bar: 2 µm. **B)** Quantification of SYP-colocalizing Homer1 cluster size and mean fluorescence intensity (MFI) in (A). (N=3, n≈300 puncta). ^ns^ non-significance, * P ≤ 0.05.

### Knockdown of TTLLs does not impair neuritic networks or neuronal function

To assess the broader impact of TTLL knockdown on neuronal networks, morphology, and function, TTLL1, TTLL4, or TTLL6 were knocked down at Day 10 and the effects on neuritic networks and dendritic branching were investigated at Day 21. Axonal and dendritic networks were studied via staining of NF-L and MAP2, respectively.

No significant changes were observed in the overall axonal (NF-L staining) or dendritic (MAP2 staining) networks across TTLL knockdown conditions compared to control (Fig. 4A-D). Sholl analysis revealed no significant differences in dendritic complexity although we noted a trend towards longer dendrites in iNeurons with TTLL1 and TTLL6 knockdowns (Fig. 4E-H). Additionally, neuronal activity, measured by microelectrode array (MEA) recordings, showed no significant alterations in spike rate or burst count across the different knockdown conditions (Fig. 4I). This indicates that iNeurons tolerate individual knockdown of the TTLLs studied here without obvious impairments in neuronal morphology and function.

**Figure 4.**
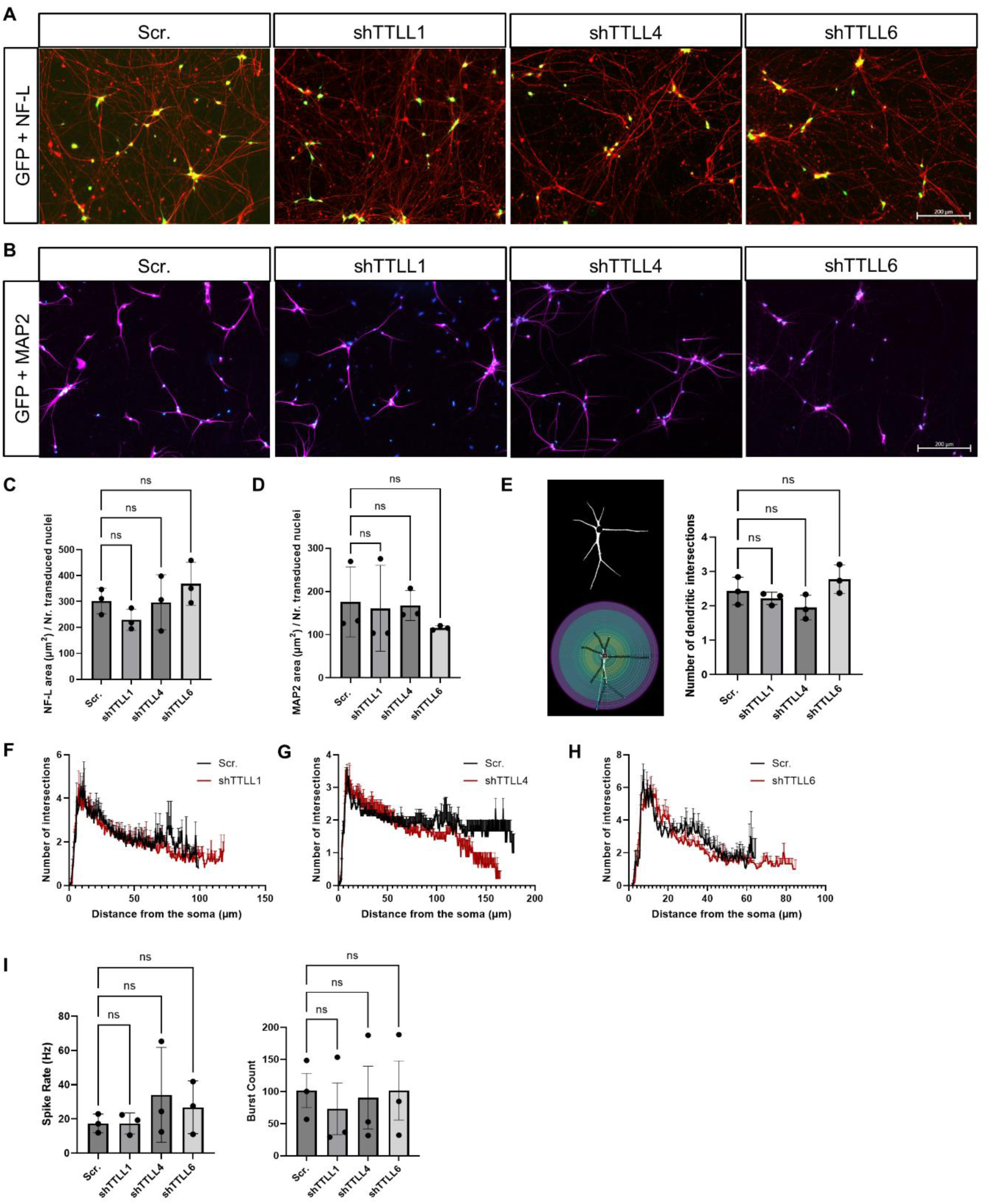
TTLL Knockdown does not affect neuronal networks, morphology, or function. A-B) Immunostaining of neurofilament-L (NF-L) (A) or MAP2 (B) in Day 21 iNeurons transduced with scrambled RNA (Scr.) or shRNAs targeting TTLL1, TTLL4, or TTLL6 for 11 days. Scale bars: 200 µm. **C)** Quantification of the area covered by the axonal network (NF-L) in (A). (N=3, n=15 fields). ^ns^ non-significance. **D)** Quantification of the area covered by the dendritic network (MAP2) in (B). (N=3, n=15 fields). ^ns^ non-significance. **E)** Quantification of the number of dendritic intersections obtained from Sholl analysis. (N=3, n=15 neurons). ^ns^ non-significance. **F-H)** Sholl profiles showing dendritic branching complexity in Day 21 iNeurons transduced with scrambled RNA (Scr.) or shRNAs targeting TTLL1 (F), TTLL4 (G), or TTLL6 (H) for 11 days. (N=3, n=15 neurons). **I)** Quantification of spike rate and burst count from microelectrode array (MEA) recordings in Day 21 iNeurons transduced with scrambled RNA (Scr.) or shRNAs targeting TTLL1, TTLL4, or TTLL6 for 11 days. (N=3, n=12-15 wells). ^ns^ non-significance.

### TTLL1 and TAU interact in HEK293T cells

To investigate the mechanistic basis behind the ameliorating effects of TTLL knockdowns against oAβ-induced toxicity and test molecular interaction and proximity, we applied Fluorescence Resonance Energy Transfer (FRET) live-cell imaging to explore potential interactions between TAU and TTLLs. TFP-TAU and YFP-TTLL1, YFP-TTLL4, or YFP-TTLL6 were co-expressed in HEK293T cells. Co-expression of TFP-TAU and YFP alone, or YFP-TTLL1, YFP-TTLL4, or YFP-TTLL6 and TFP alone, served as negative controls.

Cells co-transfected with TFP-TAU and YFP-TTLL1 showed significantly higher FRET efficiency compared to the corresponding negative controls, suggesting direct interaction between TAU and TTLL1. In contrast, no significant FRET signal was observed for TTLL4 or TTLL6 compared to their controls (Fig. 5).

**Figure 5.**
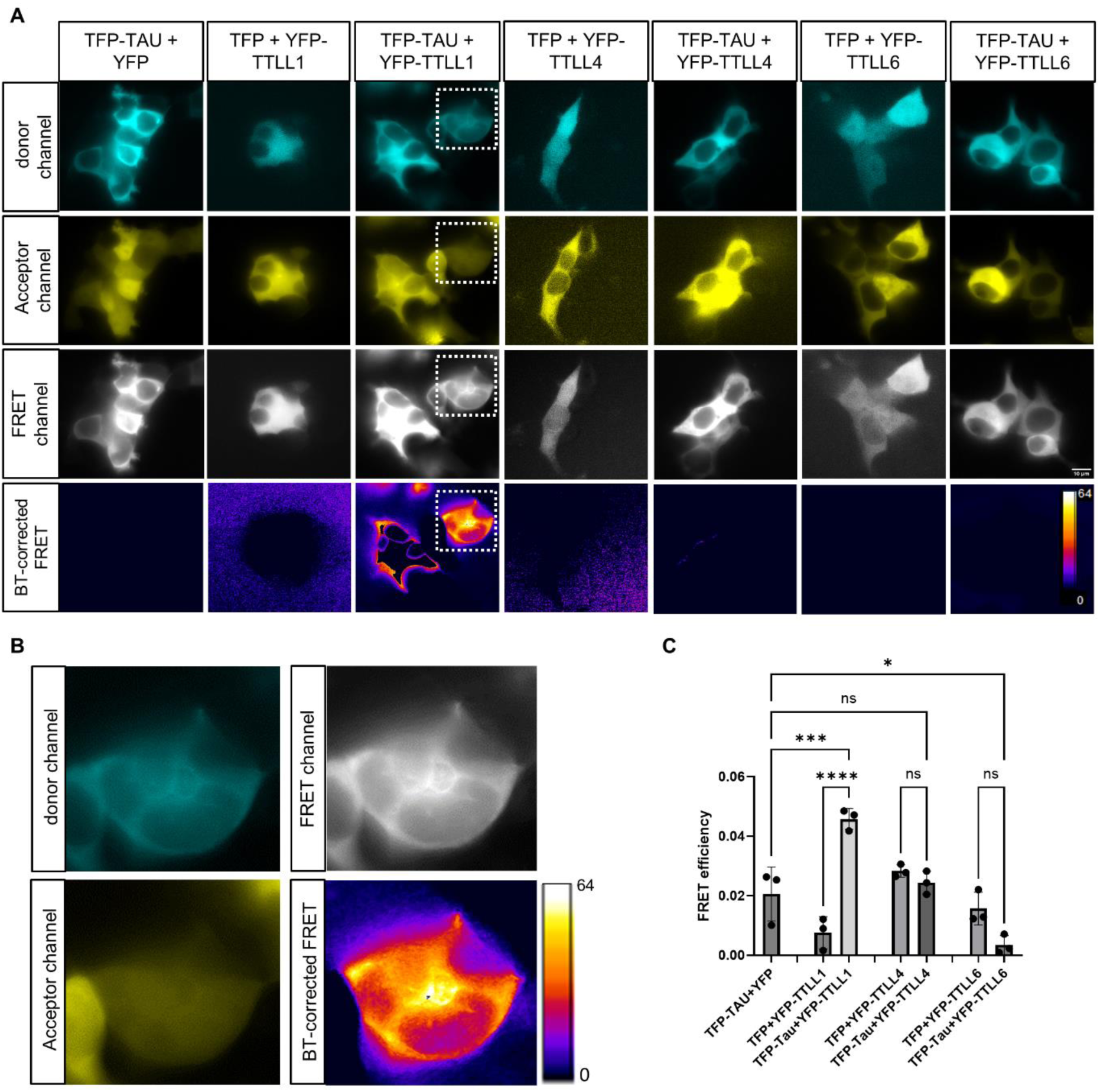
Fluorescence Resonance Energy Transfer (FRET) analysis reveals potential interaction between TAU and TTLL1. **A)** FRET microscopy images in HEK293T cells showing various controls (TFP-TAU+YFP, TFP+YFP-TTLL1, TFP+YFP-TTLL4, TFP+YFP-TTLL6) and experimental conditions (TFP-TAU+YFP-TTLL1, TFP-TAU+YFP-TTLL4, TFP-TAU+YFP-TTLL6). Images of the three detection channels: donor (TFP), acceptor (YFP), and FRET, as well as the spectral bleed-through (BT) corrected FRET are depicted. Scale bar: 10 µm. **B)** 7-fold magnification of the area framed by dashed lines in (A) showing enhanced FRET signal in low-expressing cells co-transfected with TFP-TAU and YFP-TTLL1. **C)** Quantification of FRET efficiency from the conditions shown in (A). (N=3, n=15 cells). ^ns^ non-significance, * P ≤ 0.05, *** P ≤ 0.001, **** P ≤ 0.0001.

## Discussion

This study presents a novel in vitro model of Aβ-induced tauopathy and microtubule impairments using human iPSC-derived cortical neurons. The model effectively recapitulates key features of TAU pathology seen in AD, including TAU missorting, microtubule destabilization, and synaptic defects. By exploring the role of TTLL proteins in this context, we provide new insights into the molecular mechanisms underlying Aβ-induced tauopathy and identify potential therapeutic targets.

oAβ are thought to be the upstream disease-causing agent in AD (Cline et al., 2018). Previously, acute oAβ treatment of rat-derived primary neurons led to TAU missorting, decreased acetylation and increased polyglutamylation of tubulin (Zempel et al., 2013). In this study, we showed oAβ-induced TAU missorting to the soma in our human iNeurons, a hallmark of early tauopathy (Thies and Mandelkow, 2007). This pathological shift in TAU localization was accompanied by decreased tubulin acetylation and tyrosination, and increased polyglutamylation. Microtubules are regulated by a complex set of PTMs that govern their dynamics and stability. Decreased acetylation for instance is a well-established marker of unstable microtubules (Maruta et al., 1986), while decreased tyrosination negatively affects neuronal polarity and neurite outgrowth (Erck et al., 2005). Polyglutamylation on the other hand recruits the microtubule-severing enzyme spastin (Lacroix et al., 2010), with hyperglutamylation linked to neurodegeneration and neuronal death (Berezniuk et al., 2012). In addition, we observed a significant decrease in the size of synaptophysin-juxtaposing Homer1 clusters following oAβ treatment. Homer1 is a post-synaptic density scaffold protein that is classified as an immediate early gene and which is induced by neuronal activity (Clifton et al., 2019). Synaptic Homer1 cluster disassembly was shown before to be driven by Aβ in primary rat neurons, which leads to the loss of synaptic structure and function (Roselli et al., 2009). Taken together, oAβ-insulted iNeurons exhibit detrimental changes in the PTMs of their microtubules, alongside TAU missorting and synaptic declustering, all of which is reminiscent of AD and related tauopathies.

We decided to knock down several glutamylating TTLLs in order to pinpoint the one responsible for pathological polyglutamylation, and to observe whether abolishing it would mitigate the harmful effects of oAβ. We opted to focus on TTLL1, TTLL4, and TTLL6 because of their expression levels and previously described pathological relevance: TTLL1 is the major polyglutamylating TTLL in the brain (Janke et al., 2005); TTLL1 and TTLL4 depletion mitigated neurodegeneration in pcd mice, a model that mainly exhibits adult-onset degeneration of cerebellar Purkinje neurons and selected thalamic neurons (Wu et al., 2022); and TTLL6 translocated to dendrites of primary neurons exposed to oAβ where it mediated polyglutamylation, spastin recruitment, and microtubule loss (Zempel et al., 2013). In our study, TTLL1 knockdown significantly protected iNeurons against oAβ-induced TAU missorting, reduced elevated levels of tubulin polyglutamylation, and partially alleviated the dissociation of synaptic clusters, suggesting that TTLL1 plays a critical role in the early stages of Aβ-induced tauopathy. On the other hand, overexpression of TTLL1 in iNeurons resulted in elevated levels of somatic TAU reminiscent of TAU missorting (Fig. S3), highlighting a potential toxic feedback loop between the two proteins. TTLL4 knockdown managed to decrease polyglutamylation levels but its effect on TAU missorting were less pronounced, indicating a rather secondary role for TTLL4. Interestingly, TTLL6 knockdown did not significantly affect TAU missorting but was the only knockdown that restored microtubule acetylation, hinting at a potential compensatory mechanism that stabilizes microtubules independently of TAU.

Polyglutamylating TTLLs are important enzymes that regulate neuronal microtubule organization, dynamics, and interactions with other proteins (Lacroix et al., 2010; Bodakuntla et al., 2020; Genova et al., 2023). Therefore, we speculated that knockdown of different TTLLs could impact neuronal morphology, networks, or activity, limiting the possibility of targeting TTLLs therapeutically. To investigate this further, we decided to study dendritic network and dendritic branching by staining for MAP2, a long-established dendritic marker (Caceres et al., 1984), and axonal network by staining for NF-L, an intermediate filament highly concentrated in axons and a biomarker of neuro-axonal damage (Khalil et al., 2024). However, neither neuritic networks nor dendritic branching were affected by knockdown of any of the three TTLLs investigated in this study. Additionally, burst count and spike rate of iNeurons measured by MEA recordings were also not impacted by TTLL depletion, indicating maintained neuronal activity and making TTLLs an attractive therapeutic target for reducing TAU pathology while preserving neuronal health.

In order to understand the mechanism through which TTLL1 could mediate the detrimental effects of oAβ insult and TAU missorting, we decided to investigate potential interactions between TFP-TAU and YFP-tagged constructs of the three TTLLs investigated in this study via FRET microscopy in HEK293T cells. It was reported before that TAU and TTLL6 overexpressed in HEK293T cells may interact directly (Zempel et al., 2013). In this study however, only cells expressing TFP-TAU and YFP-TTLL1 showed FRET efficiency values higher than negative controls. FRET phenomena can only occur when the distance between donor (TFP) and acceptor (YFP) is less than 10 nm, which indicates that TAU and TTLL1 are in close proximity and may interact directly with each other. This may explain why the most effective protection against oAβ was observed in iNeurons with TTLL1 knockdown. We think that missorted TAU following oAβ insult transports TTLL1 to the somatodendritic compartments where it polyglutamylates microtubules, culminating in decreased microtubule stability and synaptic loss.

## Conclusion

We showed that human iPSC-derived neurons subjected to oAβ suffer from TAU missorting, stability-decreasing changes of microtubule PTMs, and dissociation of synaptic clusters. These noxious effects are significantly reduced via TTLL1 knockdown, without affecting neuronal networks and activity. Our findings suggest that TTLL1, alone or with combination with another TTLL, could be a promising therapeutic target for preventing or slowing the progression of tauopathy in AD. Future studies should explore the therapeutic potential of genetic or pharmacological targeting of TTLL1 in vivo and investigate the broader implications of TTLL-mediated microtubule modifications in neurodegenerative diseases.

## List of Abbreviations

AD: Alzheimer disease
AUC: Area under the curve
BSA: bovine serum albumin
CCP: cytosolic carboxypeptidases
FRET: Fluorescence Resonance Energy Transfer
GFP: Green fluorescent protein
HFIP: Hexafluoro-2-propanol
HRP: horseradish peroxidase
hiPSCs: Human induced pluripotent stem cells
iNeurons: hiPSC-derived neurons
KD: Knockdown
MAP2: Microtubule-associated protein 2
MEA: Microelectrode array
MFI: Mean fluorescence intensity
NF-L: Neurofilament light chain
Ngn2: Neurogenin-2
oAβ: Oligomeric amyloid beta
Pcd: Purkinje cell degeneration
PTM: Posttranslational modification
PVDF: polyvinylidene fluoride
ROI: Region of interest
SDS: Sodium dodecyl sulfate
shRNA: short hairpin RNA
TBS-T: Tris-buffered saline with 0.1% Tween
TFP: Teal fluorescent protein
TTLL: Tubulin-Tyrosine-Ligase-Like
YFP: Yellow fluorescent protein

## Declarations

### Ethics approval and consent to participate

not applicable.

### Consent for publication

not applicable

### Availability of data and materials

The datasets used and/or analyzed during the current study are available from the corresponding author on reasonable request.

### Authors’ contributions

Study design: MAAK, HZ. Experimental work: MAAK. Methodological support: TW, JK. Data analysis and interpretation: MAAK, DA, HZ. Manuscript writing: MAAK. Manuscript commenting and proofreading: HZ, DA, TW, JK. All authors read and approved the final manuscript.

## Acknowledgments

We thank Prof. Dr. Li Gan (Weill Cornell Medicine, NY, USA) for providing Ngn2-WTC11 iPSCs. We thank Prof. Dr. Florian Klein (Institute of Virology, University Hospital Cologne) for providing lentiviral vectors. We thank Dr. Sarah Buchholz (current address: Max Planck Institute for Biology of Ageing, Cologne, Germany) for the stimulating discussions and helpful insights. Stem cell work was performed at the iPSC-Lab core facility of the CMMC (Cologne, Germany).

## References

Bachmann S, Linde J, Bell M, Spehr M, Zempel H, Zimmer-Bensch G. DNA Methyltransferase 1 (DNMT1) shapes neuronal activity of human iPSC-derived glutamatergic cortical neurons. Int J Mol Sci. 2021;22(4):2034. doi: 10.3390/ijms22042034.

Berezniuk I, Vu HT, Lyons PJ, Sironi JJ, Xiao H, Burd B, et al. Cytosolic carboxypeptidase 1 is involved in processing α- and β-tubulin. J Biol Chem. 2012;287(9):6503–17. doi: 10.1074/jbc.M111.309138.

Bodakuntla S, Schnitzler A, Villablanca C, Gonzalez-Billault C, Bieche I, Janke C, et al. Tubulin polyglutamylation is a general traffic-control mechanism in hippocampal neurons. J Cell Sci. 2020;133(3). doi: 10.1242/jcs.241802.

Buchholz S, Bell-Simons M, Cagmak C, Klimek J, Gan L, Zempel H. Cultivation, differentiation, and lentiviral transduction of human induced pluripotent stem cell (hiPSC)-derived glutamatergic neurons for studying human TAU. In: TAU protein: Methods and Protocols. Springer; 2024.

Buchholz S, Al Kabbani MA, Bell-Simons M, Kluge L, Cagmak C, Klimek J, Haag N, Iohan LC, Coulon A, Costa MR, Kilinc D, Zempel H. The tau isoform 1N4R confers vulnerability of MAPT knockout human iPSC-derived neurons to amyloid beta and phosphorylated tau-induced neuronal dysfunction. Alzheimers Dement. 2025;21(5):e14403. doi: 10.1002/alz.14403.

Caceres A, Banker G, Steward O, Binder L, Payne M. MAP2 is localized to the dendrites of hippocampal neurons which develop in culture. Brain Res. 1984;315(2):314–8. doi: 10.1016/0165-3806(84)90167-6.

Cash AD, Aliev G, Siedlak SL, Nunomura A, Fujioka H, Zhu X, et al. Microtubule reduction in Alzheimer’s disease and aging is independent of TAU filament formation. Am J Pathol. 2003;162(5):1623–7. doi: 10.1016/s0002-9440(10)64296-4.

Clifton NE, Trent S, Thomas KL, Hall J. Regulation and function of activity-dependent Homer in synaptic plasticity. Mol Neuropsychiatry. 2019;5(3):147–61. doi: 10.1159/000500267.

Cline EN, Bicca MA, Viola KL, Klein WL. The Amyloid-β oligomer hypothesis: Beginning of the third decade. J Alzheimers Dis. 2018;64(s1). doi: 10.3233/JAD-179941.

Erck C, Peris L, Andrieux A, Meissirel C, Gruber AD, Vernet M, et al. A vital role of tubulin-tyrosine-ligase for neuronal organization. Proc Natl Acad Sci U S A. 2005;102(22):7853–8. doi: 10.1073/pnas.0409626102.

Genova M, Grycova L, Puttrich V, Magiera MM, Lansky Z, Janke C, et al. Tubulin polyglutamylation differentially regulates microtubule-interacting proteins. EMBO J. 2023;42(5). doi: 10.15252/embj.2022112101.

Goodson HV, Jonasson EM. Microtubules and microtubule-associated proteins. Cold Spring Harb Perspect Biol. 2018;10(6). doi: 10.1101/cshperspect.a022608.

Hachet-Haas M, Converset N, Marchal O, Matthes H, Gioria S, Galzi JL, et al. FRET and colocalization analyzer--a method to validate measurements of sensitized emission FRET acquired by confocal microscopy and available as an ImageJ Plug-in. Microsc Res Tech. 2006;69(12):941–56. doi: 10.1002/jemt.20376.

Janke C, Rogowski K, Wloga D, Regnard C, Kajava AV, Strub JM, et al. Tubulin polyglutamylase enzymes are members of the TTL domain protein family. Science. 2005;308(5729):1758–62. doi: 10.1126/science.1113010.

Janke C, Kneussel M. Tubulin post-translational modifications: encoding functions on the neuronal microtubule cytoskeleton. Trends Neurosci. 2010;33(8):362–72. doi: 10.1016/j.tins.2010.05.001.

Jean DC, Baas PW. It cuts two ways: microtubule loss during Alzheimer disease. EMBO J. 2013;32(22):2900–2. doi: 10.1038/emboj.2013.219.

Khalil M, Teunissen CE, Lehmann S, Otto M, Piehl F, Ziemssen T, et al. Neurofilaments as biomarkers in neurological disorders - towards clinical application. Nat Rev Neurol. 2024;20(5):269–87. doi: 10.1038/s41582-024-00955-x.

Lacroix B, van Dijk J, Gold ND, Guizetti J, Aldrian-Herrada G, Rogowski K, et al. Tubulin polyglutamylation stimulates spastin-mediated microtubule severing. J Cell Biol. 2010;189(6):945–54. doi: 10.1083/jcb.201001024.

Li J, Snyder EY, Tang FH, Pasqualini R, Arap W, Sidman RL. Nna1 gene deficiency triggers Purkinje neuron death by tubulin hyperglutamylation and ER dysfunction. JCI Insight. 2020;5(19). doi: 10.1172/jci.insight.136078.

Magiera MM, Singh P, Gadadhar S, Janke C. Tubulin posttranslational modifications and emerging links to human disease. Cell. 2018;173(6):1323–7. doi: 10.1016/j.cell.2018.05.018.

Maruta H, Greer K, Rosenbaum JL. The acetylation of alpha-tubulin and its relationship to the assembly and disassembly of microtubules. J Cell Biol. 1986;103(2):571–9. doi: 10.1083/jcb.103.2.571.

Miyaoka Y, Chan AH, Judge LM, Yoo J, Huang M, Nguyen TD, et al. Isolation of single-base genome-edited human iPS cells without antibiotic selection. Nat Methods. 2014;11:291–3. doi: 10.1038/nmeth.2840.

R Core Team. R: A language and environment for statistical computing. Vienna, Austria: R Foundation for Statistical Computing; 2024. Available from: https://www.R-project.org/.

Rogowski K, van Dijk J, Magiera MM, Bosc C, Deloulme JC, Bosson A, et al. A family of protein-deglutamylating enzymes associated with neurodegeneration. Cell. 2010;143(4):564–78. doi: 10.1016/j.cell.2010.10.014.

Roselli F, Hutzler P, Wegerich Y, Livrea P, Almeida OF. Disassembly of shank and homer synaptic clusters is driven by soluble beta-amyloid(1-40) through divergent NMDAR-dependent signalling pathways. PLoS One. 2009;4(6). doi: 10.1371/journal.pone.0006011.

Sakakibara A, Ando R, Sapir T, Tanaka T. Microtubule dynamics in neuronal morphogenesis. Open Biol. 2013;3(7):130061. doi: 10.1098/rsob.130061.

Signorell A. DescTools: tools for descriptive statistics. R package version 0.99.54; 2024. Available from: https://CRAN.R-project.org/package=DescTools.

Sholl DA. Dendritic organization in the neurons of the visual and motor cortices of the cat. J Anat. 1953;87(4):387–406.

Thies E, Mandelkow EM. Missorting of TAU in neurons causes degeneration of synapses that can be rescued by the kinase MARK2/Par-1. J Neurosci. 2007;27(11):2896–907. doi: 10.1523/JNEUROSCI.4674-06.2007.

Wang C, Ward ME, Chen R, Liu K, Tracy TE, Chen X, et al. Scalable production of iPSC-derived human neurons to identify TAU-lowering compounds by high-content screening. Stem Cell Reports. 2017;9(4):1221–33. doi: 10.1016/j.stemcr.2017.08.019.

Wu HY, Rong Y, Bansal PK, Wei P, Guo H, Morgan JI. TTLL1 and TTLL4 polyglutamylases are required for the neurodegenerative phenotypes in pcd mice. PLoS Genet. 2022;18(4). doi: 10.1371/journal.pgen.1010144.

Zempel H, Thies E, Mandelkow E, Mandelkow EM. Abeta oligomers cause localized Ca(2+) elevation, missorting of endogenous TAU into dendrites, TAU phosphorylation, and destruction of microtubules and spines. J Neurosci. 2010;30(36):11938–50. doi: 10.1523/JNEUROSCI.2357-10.2010. Erratum in: J Neurosci. 32(17):6052.

Zempel H, Luedtke J, Kumar Y, Biernat J, Dawson H, Mandelkow E, et al. Amyloid-β oligomers induce synaptic damage via TAU-dependent microtubule severing by TTLL6 and spastin. EMBO J. 2013;32(22):2920–37. doi: 10.1038/emboj.2013.207.

Zempel H, Mandelkow EM. TAU missorting and spastin-induced microtubule disruption in neurodegeneration: Alzheimer disease and hereditary spastic paraplegia. Mol Neurodegener. 2015;10:68. doi: 10.1186/s13024-015-0064-1.

