## Supplemental Data for "Knockdown of TTLL1 reduces Aβ-induced TAU pathology in human iPSC-derived cortical neurons"

### Supplemental Materials

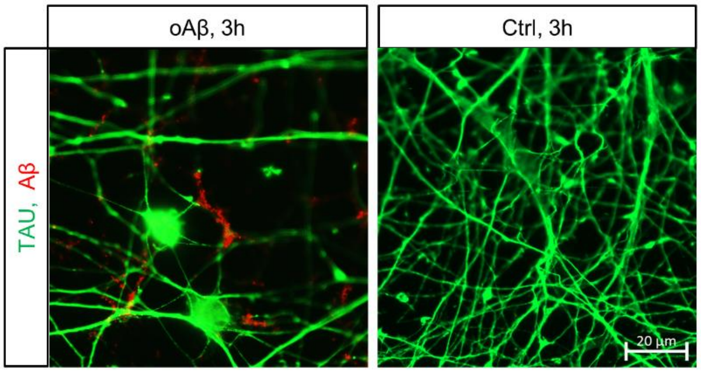


**Figure S1. oAβ localizes to neurites of iNeurons after 3 hours of treatment.** Immunostaining for TAU and Aβ in oAβ-treated iNeurons, or untreated controls (ctrl). Scale bar: 20 µm.

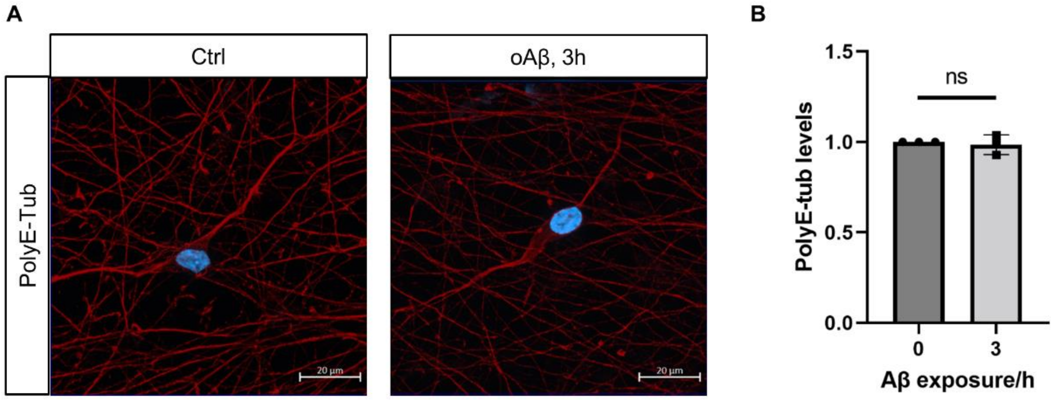


**Figure S2. *MAPT* knockout protects iNeurons against oAβ-induced elevation in tubulin polyglutamylation. A)** Immunostaining for polyglutamylated tubulin (PolyE-tub) in *MAPT* KO iNeurons treated with oAβ or untreated control (Ctrl). Scale bar: 20 µm. **B)** Quantification of somatic polyE-tub levels in (A). (N=3, n=45 neurons).

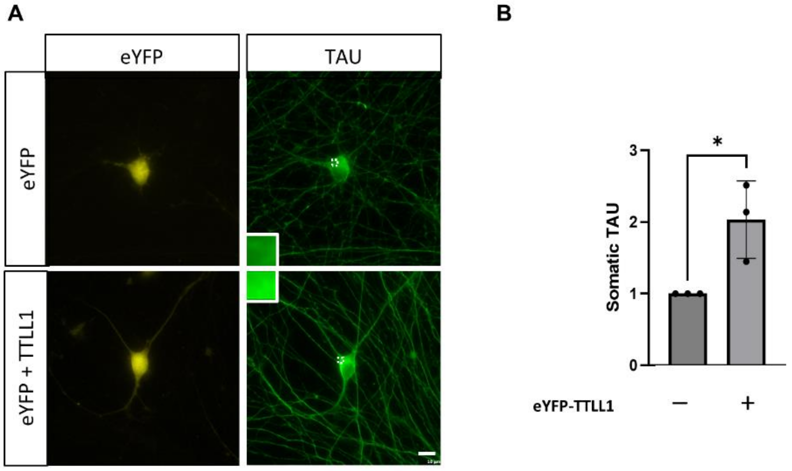


**Figure S3. TTLL1 overexpression increases somatic TAU levels in iNeurons. A)** Fluorescent eYFP signal and TAU immunostaining in Day 21 iNeurons transduced with either a transduction control or eYFP-tagged TTLL1 for 16 days. eYFP marks transduced neurons. Insets show magnification of areas framed with dashed lines, highlighting increased somatic TAU localization upon TTLL1 overexpression. Scale bar: 10 µm. **B)** Quantification of somatic TAU levels in (A). (N=3, n=45 neurons). *P ≤ 0.05.

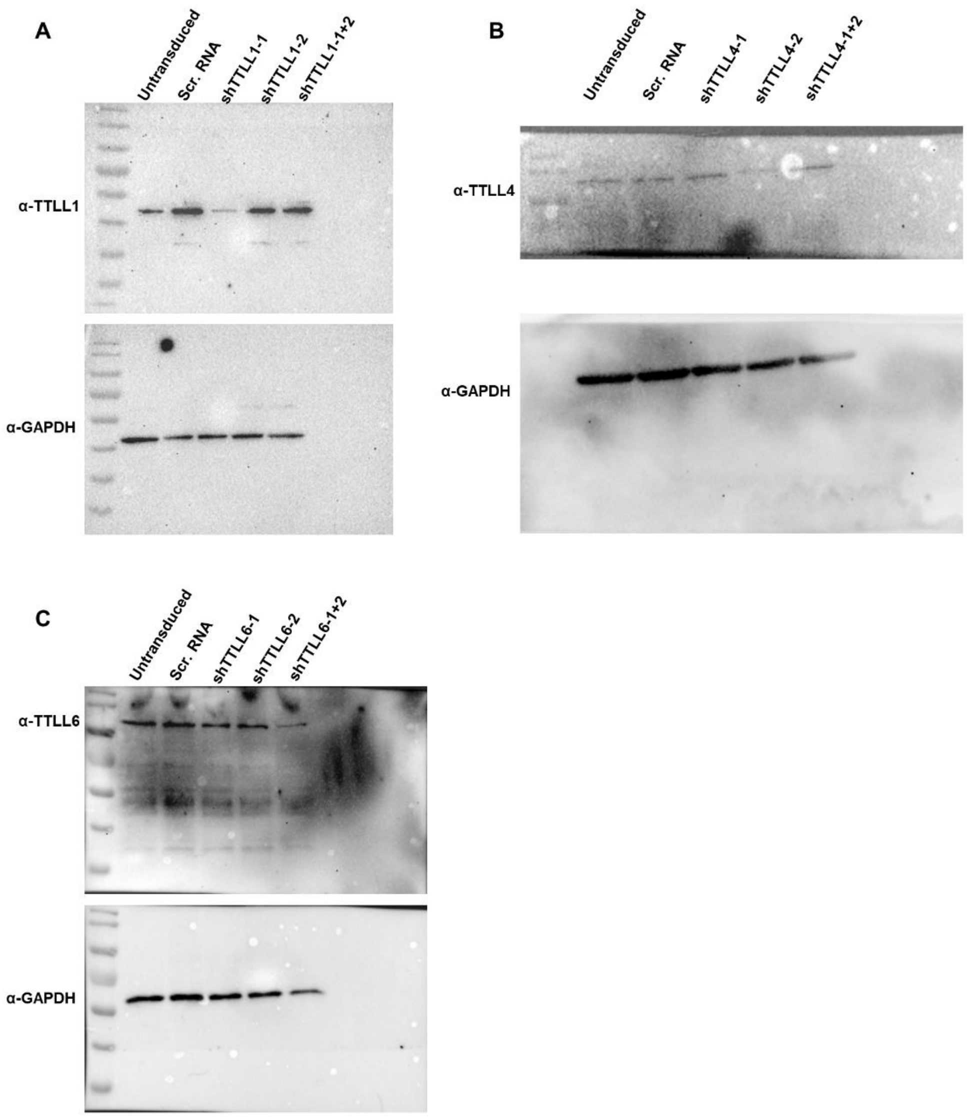


**Figure S4. Uncropped blots from Figure 2.** Immunoblotting confirmation of successful knockdown of TTLL1 (A), TTLL4 (B), and TTLL6 (C) in iNeurons.
